# LncRNA and predictive model to improve the diagnosis of clinically diagnosed pulmonary tuberculosis

**DOI:** 10.1101/863050

**Authors:** Xuejiao Hu, Hao Chen, Shun Liao, Hao Bai, Shubham Gupta, Yi Zhou, Juan Zhou, Lin Jiao, Lijuan Wu, Minjin Wang, Xuerong Chen, Yanhong Zhou, Xiaojun Lu, Tony Y Hu, Zhaolei Zhang, Binwu Ying

**Affiliations:** Department of Laboratory Medicine, West China Hospital, Sichuan University, Chengdu 610041, P. R China; Division of Laboratory Medicine, Provincial People’s Hospital, Guangdong Academy of Medical Sciences, Guangzhou 510000, P. R China; Department of Computer Science, University of Toronto, Toronto, ON, Canada; Department of Molecular Genetics, University of Toronto, Toronto, ON, Canada; The Donnelly Centre for Cellular and Biomolecular Research, University of Toronto, Toronto, ON, Canada; Department of Respiratory and Critical Care Medicine, West China Hospital, Sichuan University, Chengdu 610041, P. R China; Center for Cellular and Molecular Diagnostics, Department of Biochemistry and Molecular Biology, School of Medicine, Tulane University, New Orleans, Louisiana 70112, United States

**Author notes:** Correspondence should be addressed to Binwu Ying. Tel: [86-028-85422751]; Fax: [86-028-85422751]; Email: [ ]. Correspondence may also be addressed to Zhaolei Zhang. Tel: [1-4169460924]; Fax: [1-4169788287]; Email: [ ]. **LIST OF ABBREVIATIONS** TB: tuberculosis; PTB: pulmonary tuberculosis; MTB: *Mycobacterium tuberculosis*; TB-IGRA: interferon-gamma release assays for tuberculosis; lncRNA: long noncoding RNA; LTBI: latent TB infection; DE: differentially expressed; EHR: electronic health record; TBM: tuberculous meningitis; non-TB DC: non-TB disease control; PBMC: peripheral blood mononuclear cell; VIF: variance inflation factor; DCA: decision curve analysis; LV: lentivirus vector; BCG: Bacillus Calmette-Guerin; LDH: lactate dehydrogenase.

**Keywords:** lncRNA, electronic health record, clinically diagnosed pulmonary tuberculosis, nomogram

## Abstract

**Background:** Clinically diagnosed pulmonary tuberculosis (PTB) patients lack *Mycobacterium tuberculosis* (MTB) microbiologic evidence, and misdiagnosis or delayed diagnosis often occurs as a consequence. We investigated the potential of lncRNAs and corresponding predictive models to diagnose these patients.

**Methods:** We enrolled 1372 subjects, including clinically diagnosed PTB patients, non-TB disease controls and healthy controls, in three cohorts (Screening, Selection and Validation). Candidate lncRNAs differentially expressed in blood samples of the PTB and healthy control groups were identified by microarray and qRT-PCR in the Screening Cohort. Logistic regression models were developed using lncRNAs and/or electronic health records (EHRs) from clinically diagnosed PTB patients and non-TB disease controls in the Selection Cohort. These models were evaluated by AUC and decision curve analysis, and the optimal model was presented as a Web-based nomogram, which was evaluated in the Validation Cohort. The biological function of lncRNAs was interrogated using ELISA, lactate dehydrogenase release analysis and flow cytometry.

**Results:** Three differentially expressed lncRNAs (*ENST00000497872, n333737, n335265*) were identified. The optimal model (i.e., nomogram) incorporated these three lncRNAs and six EHR variables (age, hemoglobin, weight loss, low-grade fever, CT calcification and TB-IGRA). The nomogram showed an AUC of 0.89, sensitivity of 0.86 and specificity of 0.82 in the Validation Cohort, which demonstrated better discrimination and clinical net benefit than the EHR model. *ENST00000497872* may regulate inflammatory cytokine production, cell death and apoptosis during MTB infection.

**Conclusions:** LncRNAs and the user-friendly nomogram could facilitate the early identification of PTB cases among suspected patients with negative MTB microbiologic evidence.

## INTRODUCTION

Tuberculosis (TB) is the leading cause of death from an infectious agent ^1^, but only 56% of plmonary tuberculosis (PTB) cases reported to WHO in 2017 were bacteriologically confirmed. Thus, approximately half of all PTB cases are clinically diagnosed worldwide, and this proportion can reach 68% in China ^1^. Clinically diagnosed PTB cases are symptomatic but lack evidence of *Mycobacterium tuberculosis* (MTB) infection by smear microscopy, culture or nucleic acid amplification test ^1–3^. The diagnostic procedure for clinically diagnosed PTB is inadequate and time-consuming and often results in misdiagnosis or delayed diagnosis ^3^, leading to an increased risk of morbidity and mortality ^4^, or overtreatment ^5^. There is thus an urgent need to develop rapid and accurate strategies to diagnose PTB cases without MTB microbiologic evidence ^6, 7^. The exploration of effective host immune-response signatures represents an attractive approach for this type of assay.

Long noncoding RNAs (lncRNAs) can function as critical regulators of inflammatory responses to infection, especially for T-cell responses ^8, 9^. Increasing evidence indicates that blood lncRNA expression profiles are closely associated with TB disease ^10–12^, suggesting lncRNAs could function as potential noninvasive biomarkers for TB detection. However, previous studies have suffered from small sample size (ranging from 66 to 510) and lack independent validation.

Recent effort has focused on establishing clinical prediction rules or predictive models for TB diagnosis based on patients’ electronic health record (EHR) information ^13–16^. Such models can cost-effectively facilitate PTB diagnosis with a limited number of clinical-radiological predictors. For example, a 6-signature model from Griesel et al. (a cough lasting ≥14 days, the inability to walk unaided, a temperature > 39°C, chest radiograph assessment, hemoglobin level and white cell count) attained an AUC of 0.81 [0.80-0.82] in seriously ill HIV-infected PTB patients ^13^. However, despite these advances, current EHR models remain insufficient for precise TB diagnosis. Compelling studies have proposed that models incorporating biomarkers and EHR information attain better performance for prediction of sepsis ^17^ and abdominal aortic aneurysm ^18^. We previously reported that combining exosomal microRNAs and EHRs in the diagnosis of tuberculous meningitis (TBM) achieved AUCs of up to 0.97 versus an AUC of 0.67 obtained using EHR alone ^19^. Based on these studies, we hypothesized that combining lncRNAs with well-defined EHR predictors could be used to develop improved predictive models to identify PTB cases that lack microbiologic evidence of MTB infection.

This study was therefore performed to investigate the diagnostic potential of lncRNAs and predictive models incorporating lncRNA and EHR data for the identification of PTB cases without microbiologic MTB evidence. This study also explored the regulatory functions of lncRNA candidates during MTB infection to evaluate the biological basis for their predictive abilities.

## MATERIAL AND METHODS

### Study design

We performed this study through a four-stage approach. LncRNAs that were differentially expressed (DE) between clinically diagnosed PTB patients and healthy subjects were profiled by microarray in the Screening Step. The expression of top five lncRNAs were then analyzed in a large prospective cohort in the Selection Step of the study, which reduced the number of five lncRNAs to three based on expression difference among groups. In the Model Training Step, lncRNAs and EHRs were used to develop predictive models for clinically diagnosed PTB patients and non-tuberculosis disease control (non-TB DC) patients, and the optimal model was visualized as a nomogram. Finally, we validated lncRNAs and the nomogram in an independent prospective cohort. Functional analyses were also performed to elucidate the biological significance of lncRNAs. The study strategy is shown in Figure 1.

**Figure 1.**
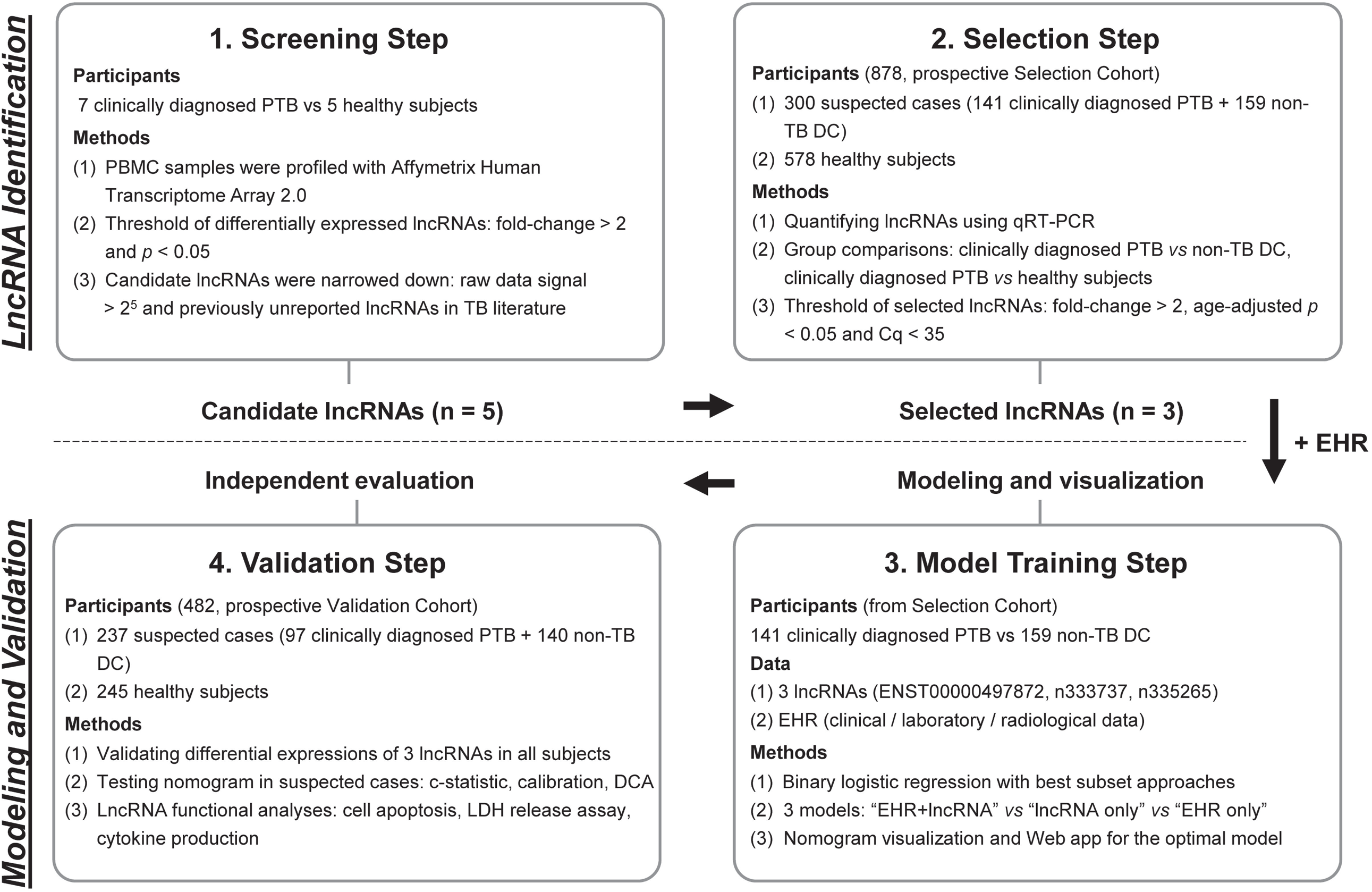
Overview of the strategy for investigating lncRNA and prediction model for clinically diagnosed PTB differential diagnosis. Abbreviations: PTB, pulmonary tuberculosis; PBMC, peripheral blood mononuclear cell; non-TB DC, non-tuberculosis disease control; DE, differentially expressed; EHR, electronic health record; DCA, decision curve analysis. LDH, lactate dehydrogenase.

### Subjects enrolment

#### Screening Cohort

We retrospectively collected age- and gender-matched 7 PTB cases and 5 healthy controls as the Screening Cohort. They were 6 males and 6 females from ages 22 to 59 years. PTB cases were clinically confirmed PTB patients with positive TB symptoms, negative MTB pathogenic examinations, and good response to anti-TB therapy. Healthy subjects had a normal physical examination and no history of TB.

#### Selection Cohort and Validation Cohort

Inpatients with clinical-radiological suspicion of PTB but lacking evidence of MTB infection were prospectively enrolled from West China Hospital between Dec 2014 and May 2017. The inclusion criteria for highly suspected patients were: (a) new patients with high clinical-radiological suspicion of PTB, (b) anti-TB therapy < 7 days on admission, (c) patients with negative MTB evidence (i.e., at least two consecutive negative smears, one negative MTB-DNA PCR and one negative culture result), (d) age ≥ 15 years, and (e) patients without severe immunosuppressive disease, HIV infection, or cardiac or renal failure. Two experienced pulmonologists reviewed and diagnosed all presumptive PTB patients, and final diagnoses for all cases were based on the combination of clinical assessment, radiological and laboratory results, response to the treatment ^1, 2^. A 12-month follow-up observation was used to confirm the classification of PTB and non-TB patients. The detailed description of patients’ symptoms and recruitment, inclusion and exclusion criteria, laboratory examinations, diagnostic criteria and procedure, treatment, and sample size estimate are provided in e-Appendix 1 and 2. In addition, healthy subjects were simultaneously recruited from a pool of healthy donors with a normal physical examination and no history of TB.

We finally enrolled a Selection Cohort of 878 participants (141 clinically diagnosed PTB, 159 non-TB DC, and 578 healthy subjects) and an independent Validation Cohort of 482 participants (97 clinically diagnosed PTB, 140 non-TB DC, and 245 healthy subjects). Details of the non-TB DC are listed in e-Table 1. Ethical approval was obtained from the Clinical Trials and Biomedical Ethics Committee of West China [no. 2014 (198)]. Informed consents were obtained from every participant.

### LncRNA detection

#### RNA isolation and cDNA preparation

Peripheral blood mononuclear cell (PBMC) samples were isolated from fresh 3 ml blood samples of each participant using a Human Lymphocyte Separation Tube Kit (Dakewe Biotech Company Limited, China). Total RNA was extracted from PBMC isolates using Trizol reagent (Invitrogen, USA). RNA concentration and purity were evaluated spectrophotometrically, and RNA integrity was determined using agarose gel electrophoresis (e-Figure 1A). The PrimeScript^™^ RT reagent Kit with gDNA Eraser (Takara, Japan) was used to remove contaminating genomic DNA and synthesize cDNA.

#### LncRNA microarray profiling

LncRNA profiles were detected using Affymetrix Human Transcriptome Array 2.0 Chips based on a standard protocol ^20^. Raw data were normalized using the Robust Multi-Array Average Expression Measure algorithm. DE lncRNAs with p-values < 0.05 and fold-changes > 2 were identified using the empirical Bayes moderated t-statistics and presented by hierarchical clustering and volcano plot ^21^. Microarray data have been deposited in the Gene Expression Omnibus under the accession GSE119143.

#### qRT-PCR for lncRNAs

LncRNA expression was measured using the SYBR^®^ Green PCR Kit (Takara, Japan) in a blinded fashion, normalized to the endogenous control *GAPDH*, and calculated according to the 2 ^−ΔΔ Cq^ method where and Cq < 35 was considered acceptable ^22^. Specific primers are presented in e-Table 2. PCR curves and the standard curve are shown in e-Figure 1B-C. Detailed methodology for RNA isolation, reverse transcription, qRT-PCR detection (procedure, quality control, product verification, and stability test) are listed in e-Appendix 3.

### Modeling

#### Data used for modeling

A total of 41 EHRs, including demographic, clinical, laboratory, and radiological findings were collected (see e-Appendix 4), and a 20% missing value threshold was applied to remove incomplete features. Features with p-values < 0.05 in univariate analysis or definite clinical significance were included for modeling. A total of 14 of the 44 original variables (41 EHRs and 3 lncRNAs) remained after filtering, including 11 EHRs and 3 lncRNAs (see e-Appendix 4).

#### Diagnostic modeling

Multivariable logistic regression was used to develop predictive models to distinguish clinically diagnosed PTB from patients with suspected PTB cases in the Selection Cohort. Feature subsets were selected and compared using the best subset selection procedure ^23^ and 10-fold cross-validation. The “EHR+lncRNA”, “lncRNA only” and “EHR only” models were developed according to their respective best feature subset in the Selection Cohort. A cutoff of each model was determined by combining the Youden’s index and the sensitivity for the samples in the training dataset equal to or greater than 0.85. The models including their cutoff were used for evaluation of the Validation Cohort.

#### Nomogram presentation and evaluation

We further adopted the nomogram to visualize the optimal model with the best AUC ^24, 25^. Nomogram calibration was assessed with the calibration curve and Hosmer-Lemeshow test (p-value > 0.05 suggested no departure from perfect fit). The performance of the nomogram was tested in the independent Validation Cohort, with total points for each patient calculated. Decision curve analysis (DCA) ^25^ was performed by evaluating the clinical net benefit of the nomogram and “EHR only” model across the overall datasets. Assessing clinical value involves comparing the nomogram and “EHR only” model using the 500 bootstrap method. The nomogram was implemented as a Web-based app using R Shiny.

### Analysis of *ENST00000497872* (lnc AL) function

The lncRNA with the most significant difference in our analysis, *ENST00000497872* (lnc AL) was analyzed in functional studies. THP-1 cells with stable overexpression and knockdown of lnc AL were constructed using recombinant lentivirus vector (LV). THP-1 cells transfected with these vectors were incubated with Bacillus Calmette-Guerin (BCG) to imitate active MTB-infection ^26^. This study examined the effect BCG exposure on THP-1 cells in five groups transfected with vectors to overexpress (LV-lnc AL) or suppress (shRNA-lnc AL) lnc AL expression, their respective empty vector constructs (LV-control and shRNA-control), or with no vector (blank control). Cell culture supernatants were harvested to measure lnc AL and the expression of six cytokines (TNF-α, IL-1β, IL-12 p70, IL-10, IFN-γ, and IL-6). Cell apoptosis and cytotoxicity after 24 h infection were detected by flow cytometry and the lactate dehydrogenase (LDH) release analysis, respectively. Detailed methodology for these experiments is presented in e-Appendix 5.

### Statistical analysis

Categorical variables were analyzed by univariate analysis with a Chi-square test and continuous variables were analyzed using Mann-Whitney U tests or Student’s t-tests. All tests were 2-sided, and p-values < 0.05 were considered statistically significant. Modeling was constructed and validated by individuals who were blinded to diagnostic categorizations. R code and data for modeling are available from https://github.com/xuejiaohu123/TBdiagnosisModel.

## RESULTS

### Characteristics of prospectively enrolled participants

The demographic and clinical characteristics of participants in the Selection and Validation Cohorts are provided in Table 1. PTB patients were younger and had greater IGRA positivity rates than their non-TB DC (p-value < 0.0001 for both the Selection and Validation Cohorts), but these groups did not differ by gender, BMI, or smoking status. Healthy subjects were age-, gender-, and BMI-matched with PTB patients, who had significantly different blood test results compared with PTB patients (Table 1).

**Table 1.** Demographic and clinical features of participants in the Selection and Validation Cohorts.

| Clinical features | Suspected clinically diagnosed PTB patients <sup>Selection</sup> |  |  | HS <sup>Selection</sup><br>(n=578) | p2 | Suspected clinically diagnosed PTB patients <sup>Validation</sup> |  |  | HS <sup>Validation</sup> (n=245) | p4 |
| --- | --- | --- | --- | --- | --- | --- | --- | --- | --- | --- |
|  | Clinically diagnosed PTB (n=141) | Non-TB DC (n=159) | p1 |  |  | Clinically diagnosed PTB (n=97) | Non-TB DC (n=140) | p3 |  |  |
| Gender, male | 84 (59.57%) | 95 (59.75%) | 0.976 | 284 (49.13%) | 0.026 | 58 (59.79%) | 87 (62.14%) | 0.715 | 126 (51.43%) | 0.162 |
| Age (years) | 37.81 ± 17.93 | 56.68 ± 14.52 | < 0.0001 | 40.59 ± 13.11 | 0.084 | 38.29 ± 17.57 | 57.96 ± 16.66 | < 0.0001 | 36.82 ± 9.28 | 0.436 |
| BMI (kg/m<sup>2</sup>) | 20.81 ± 2.99 | 20.43 ± 4.03 | 0.359 | 20.65 ± 3.19 | 0.57 | 21.59 ± 3.43 | 21.29 ± 3.62 | 0.52 | 21.51 ± 3.52 | 0.843 |
| Smoking | 61 (43.26%) | 72 (45.28%) | 0.725 | 161 (27.85%) | < 0.0001 | 41 (42.27%) | 66 (47.14%) | 0.458 | 84 (34.29%) | 0.167 |
| Radiologic pathology | 116(82.26%) | 140 (88.05%) | 0.158 | - |  | 86 (88.66%) | 130 (92.86%) | 0.264 | - |  |
| Laboratory tests |  |  |  |  |  |  |  |  |  |  |
| Positive TB-IGRA | 97 (68.88%) | 56 (35.22%) | < 0.0001 | - |  | 64 (66.00%) | 42 (30.00%) | < 0.0001 | - |  |
| C-reactive protein (mg/L) | 16.30 (5.32-54.05) | 17.80 (6.47-60.20) | 0.427 | - |  | 13.60 (3.34-43.55) | 18.60 (6.56-70.63) | 0.037 | - |  |
| Hematocrit | 0.37 ± 0.06 | 0.36 ± 0.07 | 0.162 | 0.44 ± 0.04 | < 0.0001 | 0.38 ± 0.07 | 0.35 ± 0.07 | 0.002 | 0.43 ± 0.04 | < 0.0001 |
| Erythrocytes (×10<sup>12</sup>/L) | 4.33 ± 0.72 | 4.03 ± 0.81 | 0.001 | 4.78 ± 0.46 | < 0.0001 | 4.46 ± 0.80 | 3.89 ± 0.91 | < 0.0001 | 4.80 ± 0.46 | < 0.0001 |
| Hemoglobin (g/L) | 122.57 ± 23.22 | 115.82 ± 25.20 | 0.017 | 144.46 ± 13.88 | < 0.0001 | 125.08 ± 24.25 | 113.11 ± 25.52 | < 0.0001 | 145.82 ± 13.73 | < 0.0001 |
| Platelets (×10<sup>9</sup>/L) | 238.00 (177.00-305.00) | 233.00 (149.00-299.00) | 0.171 | 193.00 (158.00-223.00) | < 0.0001 | 220.00 (160.50-313.00) | 199.50 (137.00-290.75) | 0.059 | 190.00 (165.00-230.00) | 0.001 |
| Leukocytes (×10<sup>9</sup>/L) | 6.03 (4.76-8.25) | 6.36 (4.71-9.07) | 0.488 | 5.92 (5.18-6.67) | 0.184 | 6.96 (5.06-9.14) | 5.93 (4.34-8.33) | 0.009 | 5.70 (4.91-6.55) | 0.217 |
| Lymphocytes (×10<sup>9</sup>/L) | 1.15 (0.80-1.52) | 1.28 (0.87-1.87) | 0.056 | 1.86 (1.55-2.19) | < 0.0001 | 1.29 (0.91-1.80) | 1.22 (0.86-1.62) | 0.343 | 1.85 (1.57-2.55) | < 0.0001 |
| Neutrophils (×10<sup>9</sup>/L) | 4.02 (3.23-5.93) | 4.08 (2.71-6.32) | 0.956 | 3.47 (2.87-4.09) | < 0.0001 | 4.03 (2.51-5.69) | 4.85 (3.21-6.70) | 0.023 | 3.36 (2.75-3.92) | 0.006 |
| Monocytes (×10<sup>9</sup>/L) | 0.47 (0.35-0.65) | 0.42 (0.26-0.61) | 0.015 | 0.36 (0.29-0.44) | < 0.0001 | 0.43 (0.30-0.64) | 0.46 (0.34-0.71) | 0.211 | 0.31 (0.25-0.39) | < 0.0001 |
| Alb (g/L) | 36.66 ± 6.78 | 36.60 ± 6.66 | 0.973 | 48.24 ± 2.67 | < 0.0001 | 37.33 ± 7.47 | 36.69 ± 7.22 | 0.509 | 47.06 ± 2.25 | < 0.0001 |
| Globin (g/L) | 31.69 ± 7.66 | 30.68 ± 8.10 | 0.269 | 28.92 ± 3.29 | 0.041 | 30.41 ± 7.89 | 29.73 ± 7.91 | 0.514 | 27.41 ± 3.19 | < 0.0001 |

Subscripted "Selection" or "Validation" refers to the Selection or Validation Cohort, respectively. Radiologic pathology refers to abnormal chest imaging, including at least one of the signs: polymorphic abnormality, calcification, cavity, bronchus sign, and pleural effusion. Abbreviations: non-TB DC, non-tuberculosis disease control patients; HS, healthy subjects. p1, p-value for the comparison of clinically diagnosed PTB patients and non-TB DCs in the Selection Cohort; p2, p-value for the comparison of clinically diagnosed PTB patients and healthy subjects in the Selection Cohort; p3, p-value for the comparison of clinically diagnosed PTB patients and non-TB DCs in the Validation Cohort; p4, p-value for the comparison of clinically diagnosed PTB patients and healthy subjects in the Validation Cohort.

Clinically diagnosed PTB patients were responsible for 29.82% (238/798) of all active PTB patients (see e-Appendix 1). This rate is markedly lower than a nationwide estimate of 68% based on primary public health institutions ^1^, but represents the clinically diagnosed PTB rate in a referral hospital with experienced specialists.

### LncRNAs microarray profiles and candidate selection

In the Screening Step, microarray profiling identified a total of 325 lncRNAs that were differentially expressed (287 upregulated and 38 downregulated) in the clinically diagnosed PTB patients versus healthy subjects. Hierarchical clustering and a volcano plot revealed clearly distinguishable lncRNA expression profiles (e-Figure 2). Top five lncRNA candidates were chosen based on a set of combined criteria: fold-change > 2 between groups, p-value < 0.05, signal intensity > 25 ^27^, and including unreported lncRNAs in TB literature ^28^. Three of these five lncRNAs were upregulated (*n335265, ENST00000518552* and *TCONS_00013664*) and two were downregulated (*n333737* and *ENST00000497872*) in PTB versus control subjects (e-Table 3).

### Differentially expressed lncRNAs in clinically diagnosed PTB

The expression level of these five candidate lncRNAs was measured by qRT-PCR in the Selection Cohort, which consisted of 141 clinically diagnosed PTB, 159 non-TB DC, and 578 healthy subjects. Two lncRNAs (*ENST00000518552* and *TCONS_00013664*) were excluded from further analysis due to their low abundance expression (Cq > 35) in this cohort. Of the three remaining lncRNAs, *ENST00000497872* and *n333737* were downregulated and *n335265* was upregulated in PTB patients versus healthy subjects (e-Table 4). Comparison between clinically diagnosed PTB cases and non-TB DC patients revealed a decreased expression of *ENST00000497872* and *n333737* in PTB patients (e-Figure 3A), age-adjusted p-values both < 0.0001).

Short-term stability, an essential prerequisite of a potential lncRNA biomarker, was assessed in PBMC samples. This study found that incubation up to 24 h had minimal effect on the expression of *ENST00000497872, n333737*, and *n335265* (e-Table 5), in accordance with a previous report of lncRNA stability in blood ^29^.

### Diagnostic modeling and nomogram visualization

Three logistic regression models, “EHR+lncRNA”, “EHR only”, and “lncRNA only” were evaluated as part of the training step in the Selection Cohort (see e-Appendix 4). The variance inflation factors between the features ranged from 1.02 to 1.29, indicating no collinearity within models. The “EHR+lncRNA” model yielded the highest AUC (0.92) for distinguishing clinically diagnosed PTB from suspected PTB patients, compared to AUCs of 0.87 and 0.82 for the “EHR only” and “lncRNA only” models, respectively (Figure 2A). The “EHR+lncRNA” model also had the best performance in sensitivity, specificity, accuracy, positive predictive value, and negative predictive value (Table 2).

**Table 2.**
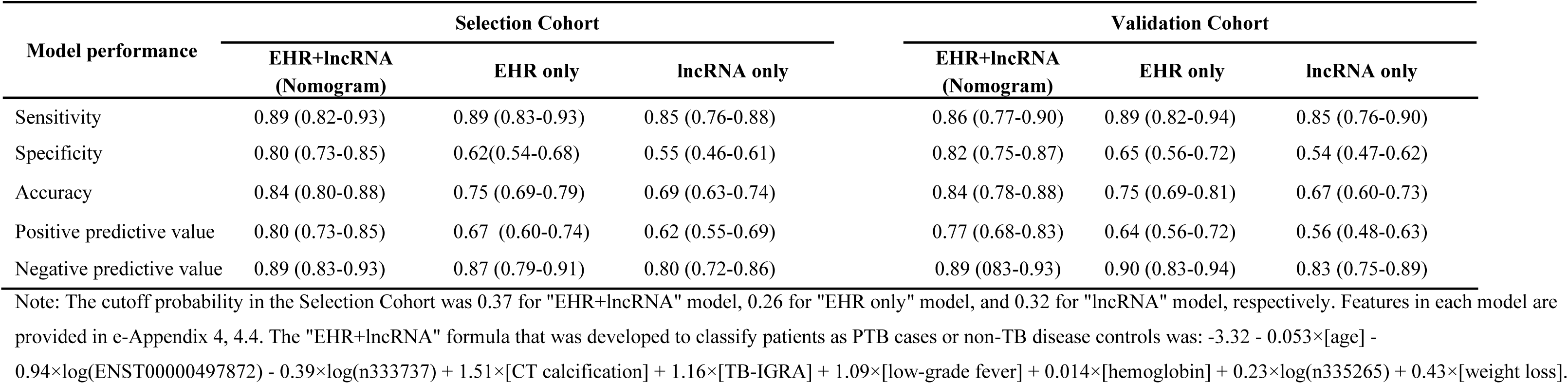
Performances of the comparative diagnostic models.

**Figure 2.**
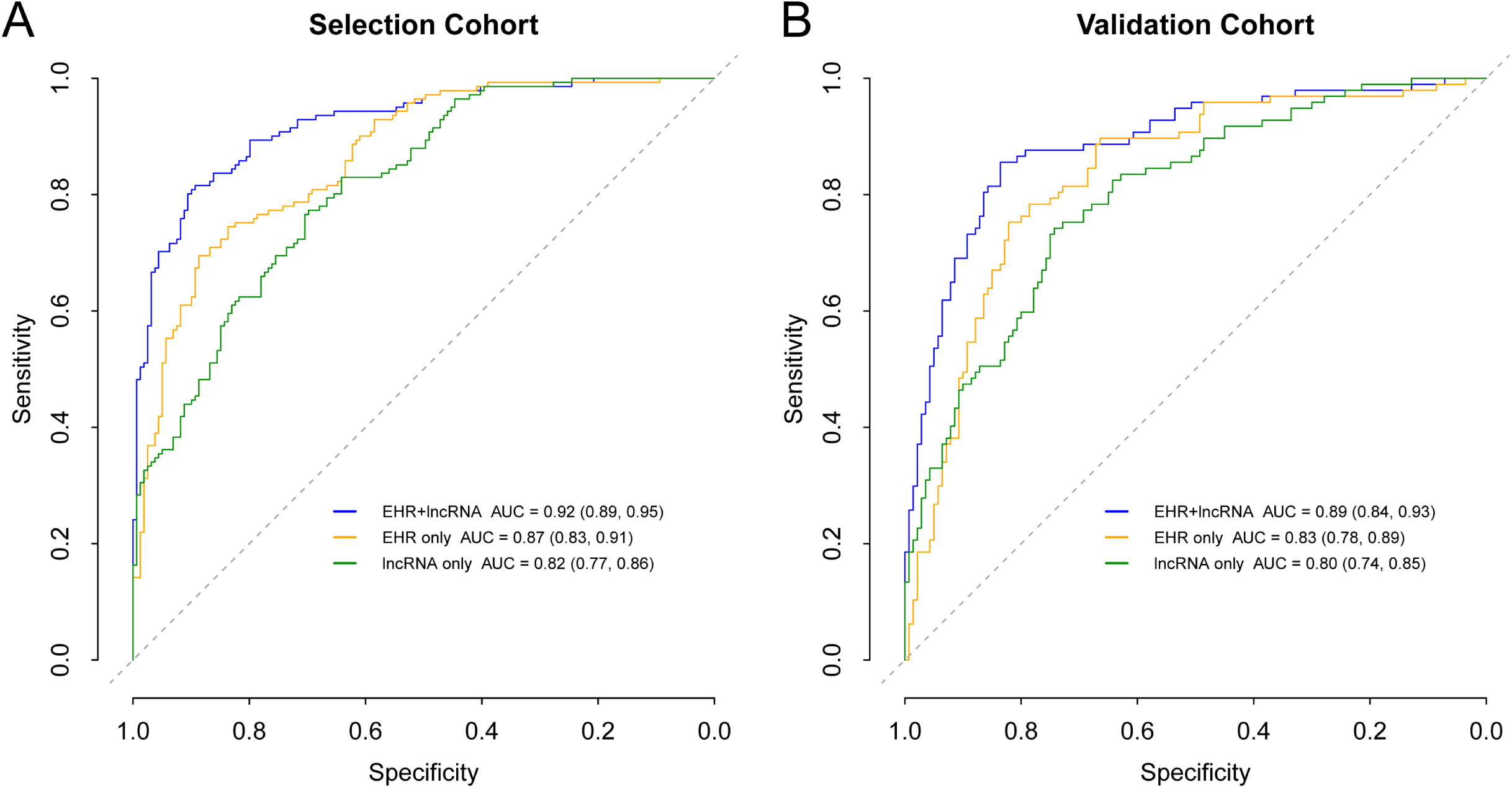
Receiver operator curves of different models in the Selection and Validation Cohort. (A), ROC of the Selction Cohort. The 10-fold cross-validation ROC of “EHR+lncRNA” model is provided in the e-Figure 5. P-values for model AUC comparisons in the Selection Cohort: 0.00012 (“EHR+lncRNA” vs “EHR only”), 1.402×10^-7^ (“EHR+lncRNA” vs “lncRNA only”), and 0.103 (“EHR only” vs “lncRNA only”), respectively. P-values < 0.016 (0.05/3) were considered statistically significant. (B), ROC of the Validation Cohort. P-values for model AUC comparisons in the Validation Cohort: 0.004 (“EHR+lncRNA” vs “EHR only”), 0.0003 (“EHR+lncRNA” vs “lncRNA only”), and 0.361 (“EHR only” vs “lncRNA only”), respectively.

The optimal “EHR+lncRNA” model was displayed as a nomogram (Figure 3A), and the top five features of the nomogram were *ENST00000497872*, age, *n333737*, CT calcification, and TB-IGRA results (e-Table 6). Seneitivity and specificity of the nomogram for prediction of clinically diagnosed PTB was 0.89 (0.82-0.93) and 0.80 (0.73-0.85) at a cutoff of 0.37 (Table 2). A calibration curve in the Selection Cohort (Figure 3B) indicated a good agreement between nomogram prediction and actual PTB cases and was confirmed by the nonsignificant Hosmer-Lemeshow test (p-value = 0.957). This nomogram was generated as a free online app (available at https://xuejiao.shinyapps.io/shiny/) to facilitate its access for other studies. This app allows the user to insert the values of specific predictors and provides the risk prediction as a whole number percentage.

**Figure 3.**
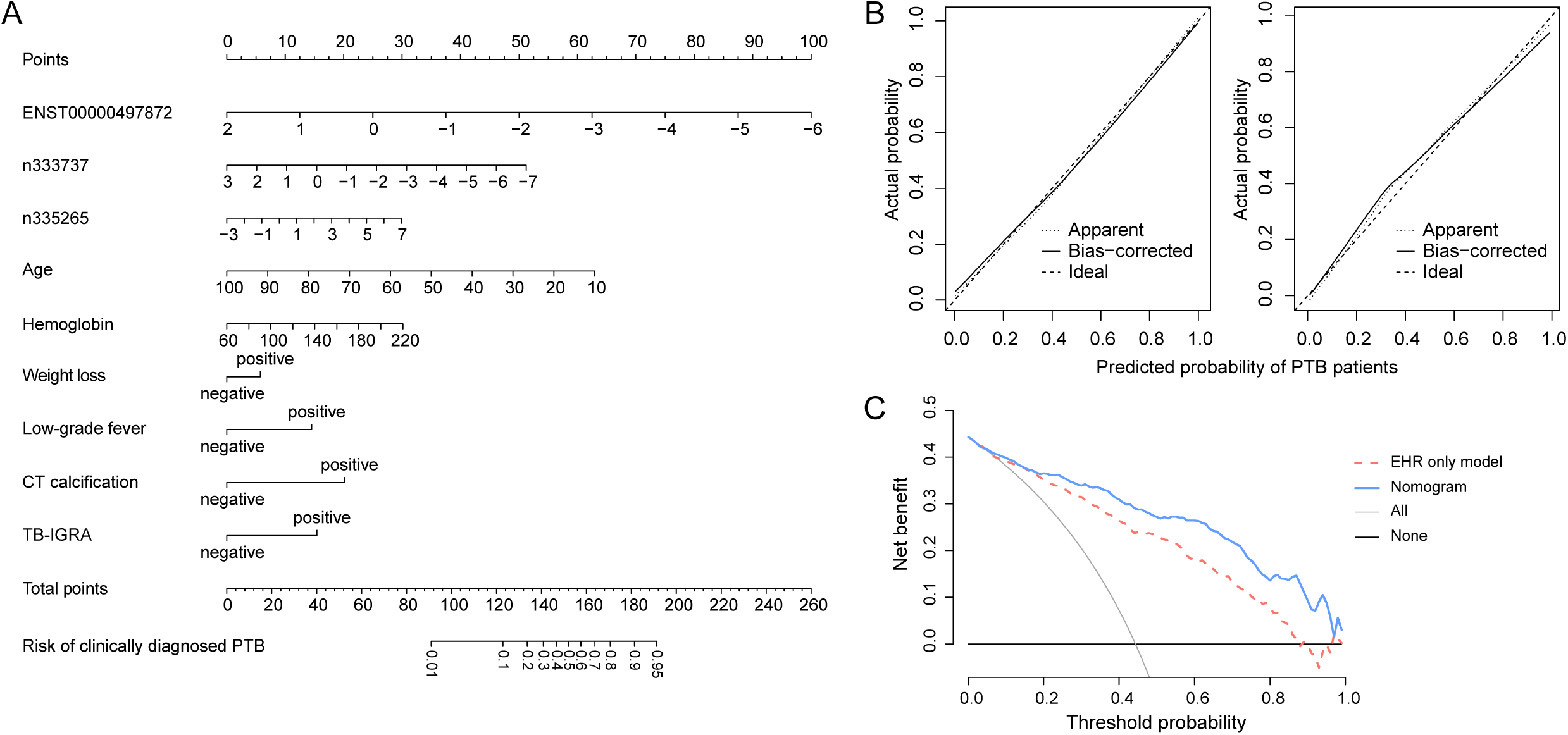
Nomogram for the prediction of clinically diagnosed PTB based on the optimal models. (A), Nomogram to predict the risk of clinically diagnosed PTB patients, in which points were assigned based on the feature rank order of the effect estimates. A vertical line is drawn between the “Point” axis and the corresponding point for each feature to generate a total point score and PTB probability. (B), Calibration plot in the Selection Cohort (left in B) and Validation Cohort (right in B), with lines indicating the ideal (dashed), apparent (dotted) and bias-corrected (unbroken) predictions of the nomogram. (C), Decision curve analysis for the nomogram and “EHR only” model with lines indicating the nomogram (blue), “EHR only” model (red dash), and assumptions that no patients or all patients have PTB (black and grey, respectively).

### Validation for lncRNAs and the nomogram

In the Validation Step, the three candidate lncRNAs were analyzed in an independent Validation Cohort contains 97 clinically diagnosed PTB cases, 140 non-TB DC and 245 healthy subjects. This analysis observed an lncRNA expression pattern similar to that observed in the Selection Cohort (e-Table 4, e-Figure 3B). All three models were applied to the Validation Cohort, and as reported in Table 2 and Figure 2 it was found that the nomogram achieved superior discrimination (AUC: 0.89 [0.84-0.93]), good calibration (Figure 3B, and p-value = 0.668 for Hosmer-Lemeshow test) for clinically diagnosed PTB prediction. The sensitivity and specificity of the nomogram at the cutoff of 0.37 in the Validation Cohort was 0.86 (0.77-0.90) and 0.82 (0.75-0.87), respectively. DCA indicated that the nomogram outperformed the conventional “EHR only” model with a higher clinical net benefit within a threshold probability range from 0.2 to 1 (Figure 3C).

### LncRNA response to anti-TB treatment

LncRNAs were next analyzed for the ability to predict anti-TB treatment response. Paired samples were collected from 22 clinically diagnosed PTB patients before and after 2-month intensive therapy ^30^, and the expressions of *ENST00000497872, n333737*, and *n335265* were measured by qRT-PCR. All these patients had good response to therapy based on the clinical and radiological findings, and *ENST00000497872* and *n333737* levels significantly increased post-treatment (p-values = 0.005 and 0.0005, respectively, Figure 4), suggesting that lncRNA expression increased in response to therapy.

**Figure 4.**
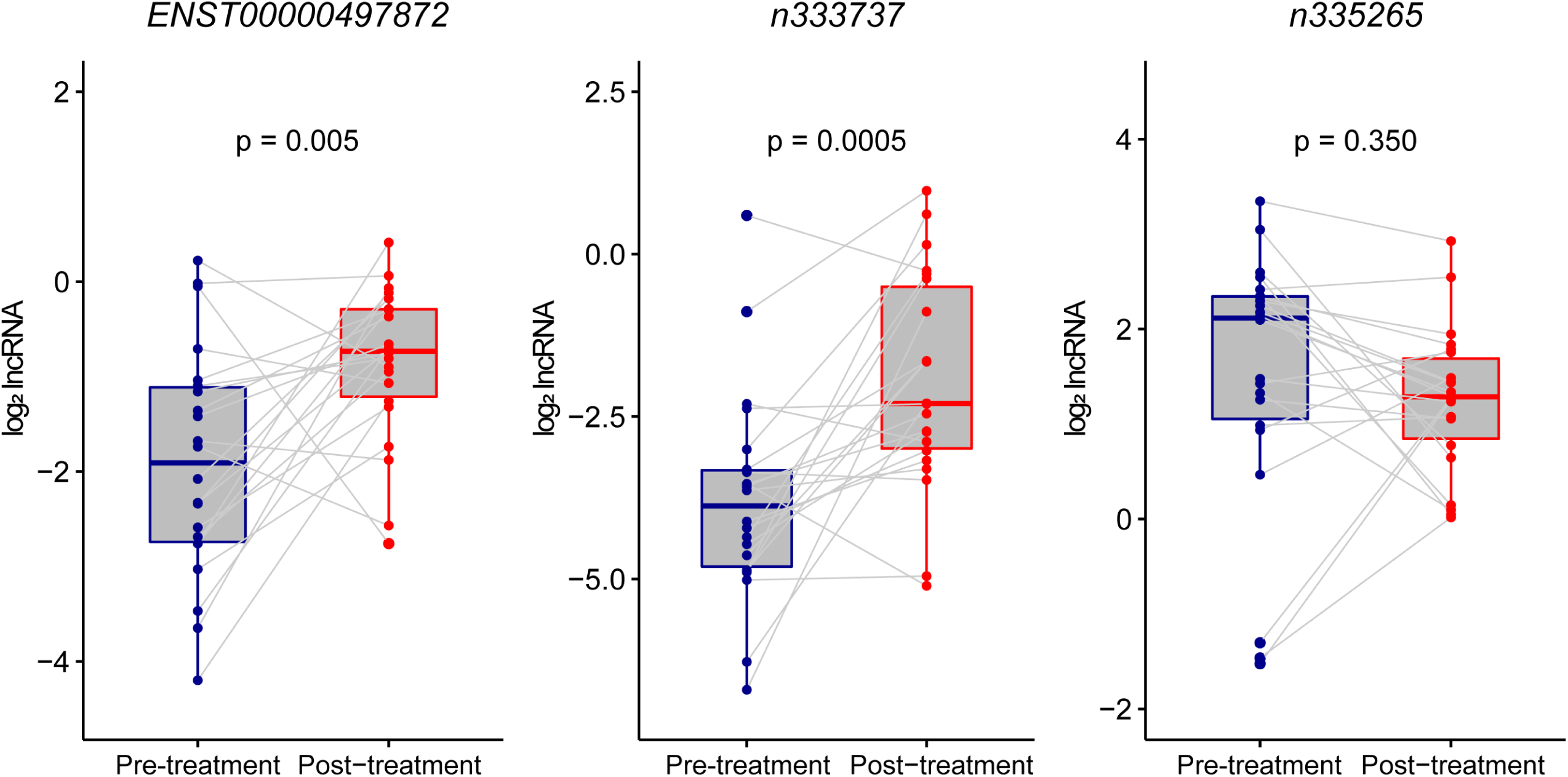
Alteration of lncRNAs before and after 2-month intensive therapy. LncRNA expressions before (blue) and after (red) a 2-month intensive anti-TB treatment regimen. Altered lncRNA expressions were calculated using log2 lncRNA (post-treatment expression / pre-treatment expression) and the Wilcoxon matched-paired rank test was used for comparisons among 22 paired samples. The median and interquartile range of log2 lncRNA were as follows: *ENST00000497872* (before: −1.91 [−2.74, −1.11]; after: −1.55 [−2.61, −0.79]), *n333737*: (before: −3.88 [−4.81, −3.33]; after: −2.30 [−2.99, −0.50]), *n335265* (before: 2.12 [1.05, 2.34]; after: 1.29 [0.85, 1.69]), respectively.

### Functional studies of *ENST00000497872*

We investigated whether *ENST00000497872* (i.e., lnc AL) could affect the host immune response. At 24 h and 48 h post BCG-infection, lnc AL overexpression (e-Figure 4) led to decreased production of proinflammatory cytokines TNF-α and IL-1β and an increase in INF-γ (Figure 5A). Conversely, knockdown of lnc AL resulted in a significant TNF-α and IL-1β increases and an INF-γ reduction. Lnc AL knockdown was also associated with an increasing trend of cell apoptosis (Figure 5B and 5C) and cell death (Figure 5D). These results implicate an inflammatory regulation of lnc AL during MTB-infection.

**Figure 5.**
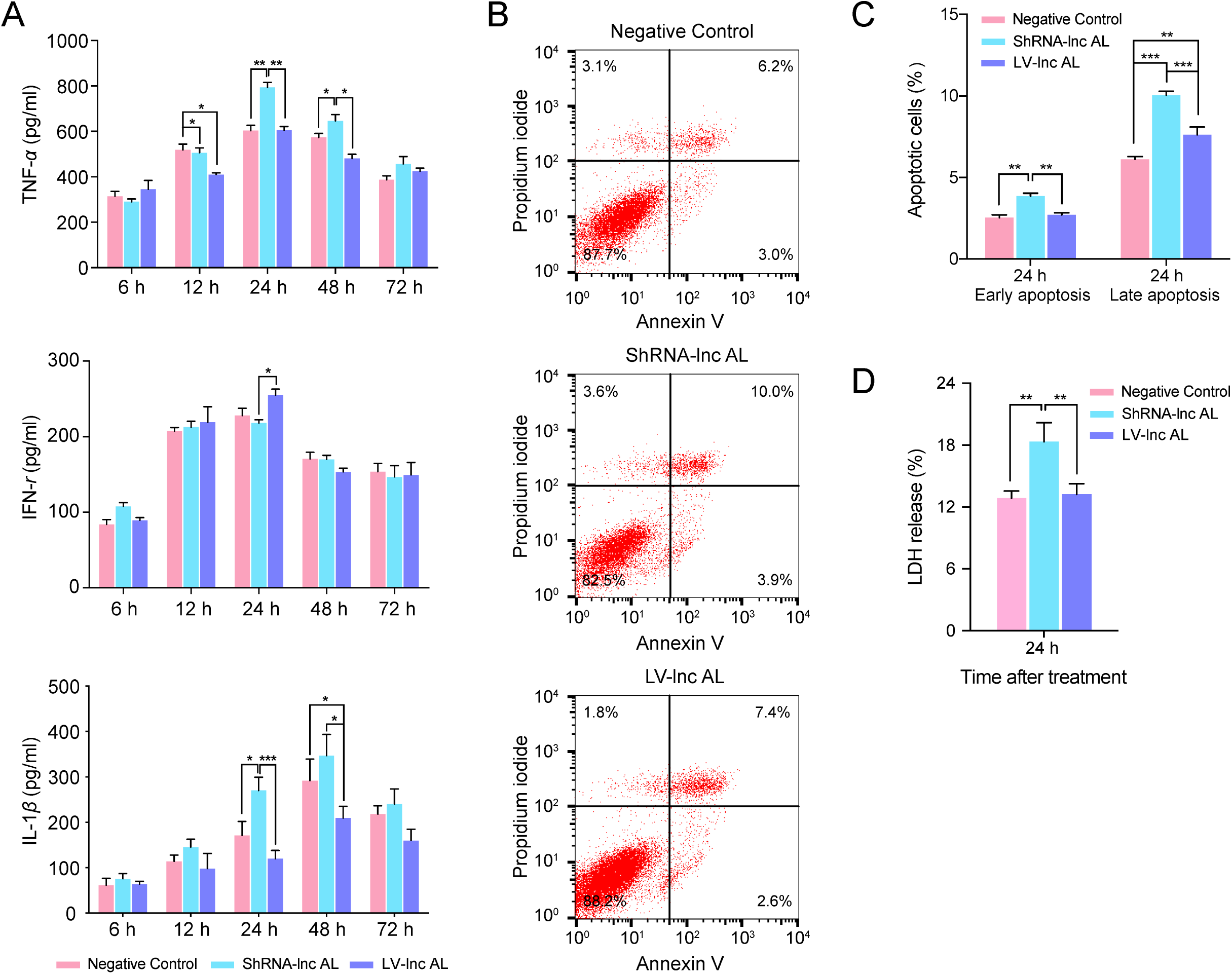
Regulation of lncRNA on inflammatory cytokine, cell apoptosis and cytotoxicity in BCG-infected THP-1 cells. (A), Cytokine expression. (B), Flow cytometry analysis of cell apoptosis. (C), Graph of apoptosis data. (D), LDH release analysis of cell cytotoxicity for BCG-infected THP-1 cells. LV-control and shRNA-control mean values considered negative control values, and the blank control is not shown. Three cytokines (IL-12 p70, IL-10 and IL-6) did not significantly differ and are not shown. Difference between groups were analyzed by one-way ANOVA and Bonferroni’s post-test comparison among groups (*p-value < 0.05, **p-value < 0.01, and ***p-value < 0.001).

## DISCUSSION

The present work focused on the challenge of accurately diagnosing PTB patients without microbiological evidence of MTB infection. Our study showed that three lncRNAs (*ENST00000497872, n333737*, and *n335265*) were potential biomarkers for clinically diagnosed PTB patients. Addition of three lncRNAs (*ENST00000497872, n333737* and *n335265*) to a conventional EHR model improved its ability to identify PTB cases from TB suspects, with the AUCs increasing from 0.83 to 0.89. The lncRNA that was most significantly enriched in the PTB group of this study, *ENST00000497872* (chr14:105703964-105704602), is located close to *IGHA1* (chr14: 105703995-105708665), and functional analyses indicated that expression of this lncRNA was involved in the regulation of inflammatory cytokine production and cell apoptosis in MTB-infected macrophages, although further studies are needed to investigate the mechanisms responsible. Consistent with published lncRNA data ^8–12, 31^, this data provide new evidence that lncRNAs could participate in TB immunoregulation and serve as promising biomarkers for TB diagnosis.

In addition to the three lncRNAs, we identified six EHR predictors (age, CT calcification, positive TB-IGRA, low-grade fever, elevated hemoglobin, and weight loss) that were essential in TB case finding, as proposed by prior findings ^15, 16^. Age was an important negative predictor for clinically diagnosed PTB, which appears to conflict with the consensus that advanced age correlates with higher TB susceptibility ^32^. This may be explained by differences in the enrollment of the PTB patients and control subjects. Previous studies included healthy and/or vulnerable subjects as controls, while we enrolled inpatients with a wide range of pulmonary diseases and older ages as disease controls.

This study serves as a first proof-of-concept study to show that integrating lncRNA signatures and EHR data could be a more promising diagnostic approach for PTB patients with negative MTB pathogenic evidence. The “EHR+lncRNA” model had good discrimination (through AUC and diagnostic parameters), reliable calibration (via calibration curve and Hosmer-Lemeshow test), and potential clinical utility for decision-making (using DCA). The “EHR+lncRNA” model avoided some common problems associated with sputum-based features, such as poor sputum quality or problematic sampling ^33^, to improve its reliability and clinical utility. Nomogram has been shown to remarkably promote early diagnosis of intestinal tuberculosis ^24^ and prognosis prediction in PTB ^34^ and TBM ^35^. “EHR+lncRNA” model herein was visualized as a nomogram and further implemented in an app. The online nomogram uses readily obtainable predictors and automatically outputs a personalized quantitative risk estimate for PTB. Utilizing this user-friendly tool may facilitate the rapid identification of PTB cases among suspected TB patients without MTB microbiologic evidence to improve TB diagnosis, especially in resource-constrained areas with high TB prevalence.

Our study has several limitations. Modeling in this study was conducted based on data from a single large hospital, and multi-center validation studies are needed. Further, because Xpert MTB/RIF is still not routinely available in most clinical laboratories of China, and since previous Xpert studies reported moderate sensitivities ranged from 28% to 73% ^36–38^ in smear-negative PTB patients, we did not consider Xpert in our research, which may limit the generalization of our findings.

In summary, a novel nomogram we developed and validated in this study that incorporated three lncRNAs and six EHR fields may be a useful predictive tool in identifying PTB patients with negative MTB pathogenic evidence, and merits further investigation.

## AUTHOR’S CONTRIBUTIONS

XH, HC, ZZ, and BY developed the concept and experimental design; XH, HB, YZ, JZ, LJ, LW, and MW enrolled patients and performed the experiments; XH, SL, and SG developed and validated the diagnostic models; XC, YZ, XL, and TYH provided expert advice and support; all authors contributed to write or revise the manuscript, provided intellectual input and gave final approval.

## ACKNOWLEDGEMENTS

We thank Dr. Yi Zhang from Gminix (Shanghai, China) for assistance with bioinformatic analyses.

## SUPPLEMENRARY FIGURE CAPTIONS

e-Figure 1. RNA electrophoresis, amplification curve of qRT-PCR and standard curve of control cDNA

e-Figure 2. Hierarchical clustering and volcano plot for differentially expressed lncRNA profiles in the Screening Cohort

e-Figure 3. LncRNA expression between clinically diagnosed PTB patients and non-TB disease controls in the Selection and Validation Cohorts

e-Figure 4. qPCR analysis of *ENST00000497872* expression in BCG-infected THP-1 cells

e-Figure 5. Ten-fold cross-validation ROC of “EHR+lncRNA” model developed using the data from the Selection Cohort

## SUPPLEMENRARY TABLE CAPTIONS

e-Table 1. Disease controls in the present study

e-Table 2. Specific qRT-PCR primers for lncRNAs

e-Table 3. Expression of five candidate lncRNAs in the Screening Cohort

e-Table 4. Comparison of lncRNA expression between clinically diagnosed PTB patients and healthy subjects in the Selection and Validation Cohorts

e-Table 5. Short-term stability evaluation of lncRNAs in PBMC samples

e-Table 6. Details of “EHR+lnRNA” logistic regression model to differentiate clinically diagnosed PTB among 300 highly suspected patients

